# How squirrels protect their caches: Location, conspicuousness during caching, and proximity to kin influence cache lifespan

**DOI:** 10.1101/738237

**Authors:** Mikel M. Delgado, Lucia F. Jacobs

## Abstract

Scatter-hoarding animals cannot physically protect individual caches, and instead utilize several behavioral strategies that are hypothesized to offer protection for caches. We validated the use of physically altered, cacheable food items, and determined that intraspecific pilfering among free-ranging fox squirrels (*N* = 23) could be assessed in the field. In this study we were able to identify specific individual squirrels who pilfered or moved caches that had been stored by a conspecific. We identified a high level of pilfering (25%) among this population. In a subsequent study, we assessed the fate of squirrel-made caches. Nineteen fox squirrels cached 294 hazelnuts with passive integrated transponder tags implanted in them. Variables collected included assessment and cache investment and protection behaviors; cache location, substrate, and conspicuousness of each cache; how long each cache remained in its original location, and the location where the cache was finally consumed. We also examined whether assessment or cache protection behaviors were related to the outcomes of buried nuts. Finally, we measured the population dynamics and heterogeneity of squirrels in this study, testing the hypothesis that cache proximity and pilferage tolerance could serve as a form of kin selection. Polymer chain reaction (PCR) was used to analyze hair samples and determine relatedness among 15 squirrels, and the potential impact of relatedness on caching behavior. Results suggested that cache protection behaviors and the lifespan of a cache were dependent on the conspicuousness of a cache. Squirrels may mitigate some of the costs of pilfering by caching closer to the caches of related squirrels than to those of non-related squirrels.

## Introduction

Scatter-hoarding animals cannot physically protect individual caches, and instead utilize several behavioral strategies that are hypothesized to offer protection for caches. These behaviors include assessing food items to appropriate allocate cache effort (e.g., Preston & Jacobs, 2009), caching out of sight of conspecifics (e.g., Dally, Emery, & Clayton, 2004), caching food items at low density (e.g., Male & Smulders, 2007), or at a great distance from the food source (Vander Wall, 1995a), or spending more time carefully covering caches (e.g., Leaver, Hopewell, Caldwell, & Mallarky, 2007). How these behaviors contribute to the survival and retrieval of these caches or might reduce pilferage from conspecifics is still unknown. In fact, little is known about what factors do contribute to whether a cache is stolen, forgotten, or retrieved by the animal who cached it.

Many behavioral mechanisms that scatter-hoarding animals could use to protect caches have yet to be examined in detail, such as the adaptive use of food assessment. Several animal species display food assessment behaviors including squirrels, primates, birds and fish (Jablonski, Fuszara, Fuszara, Jeong, & Lee, 2015; Kislalioglu & Gibson, 1976; Melin et al., 2009; Preston & Jacobs, 2009). These behaviors help animals select higher quality food items, as demonstrated in scatter-hoarding Western scrub jays (*Aphelocoma californica*) and Piñon jays (*Gymnorhinus cyanocephalus*), who use bill clicking and item handling to choose heavier seeds (Langen & Gibson, 1998; Ligon & Martin, 1974).

In the case of food-storing animals, assessment may provide information that allows for the adjustment of cache investments to the value of individual food items. Fox squirrels (*Sciurus niger*) use two overt behaviors to assess food items, head flicks and paw manipulations. These behaviors may help squirrels assess the quality, weight, and perishability of food items before caching or eating them (Delgado, Nicholas, Petrie, & Jacobs, 2014; Preston & Jacobs, 2009). For example, fox squirrels are significantly more likely to cache than eat items after they perform a head flick (Delgado et al., 2014; Preston & Jacobs, 2009). Because many scatter-hoarding animals, including squirrels, jays, mice, and chipmunks, adjust cache distance to the value of food (e.g., Delgado et al., 2014; Jokinen & Suhonen, 1995; Moore, McEuen, Swihart, Contreras, & Steele, 2007; Tamura, Hashimoto, & Hayashi,1999; Waite & Reeve, 1995), it follows that they should have some means of assessing individual food items to determine their value.

Several scatter-hoarding animals, including squirrels, are sensitive to the presence of other animals and adjust caching behaviors when competitors are present (Dally, Clayton, & Emery, 2006; Dally, Emery, & Clayton, 2005; Emery, Dally, & Clayton, 2004). Birds in the corvid and parid families eat food items and reduce the number they cache, or wait to cache until after competitors have left (Goodwin, 1956; James & Verbeek, 1984; Lahti & Rytkönen, 1996; Leaver et al., 2007; Stone & Baker, 1989). Western scrub jays cache out of view or move their caches several times when conspecifics are present, presumably to reduce visual cues available to competitors (Dally et al., 2004; Dally et al., 2005). Eurasian jays (*Garrulus glandarius*) may even reduce acoustic information available to competitors by caching in quieter substrate (Shaw & Clayton, 2013), as other jays appear to use auditory information to locate and steal caches made by other jays (Shaw & Clayton, 2014). Scatter-hoarding tree squirrels also vary several behaviors in the presence of competitors: the amount of time and effort spent traveling to a cache site (Delgado et al., 2014; Hopewell, Leaver, & Lea, 2008; Leaver et al., 2007), the number of holes dug before selecting a final cache location (Delgado et al., 2014; Steele et al., 2008), and time spent covering a cache site with available substrate such as dirt or leaves (Delgado et al., 2014; Hopewell & Leaver, 2008).

These behaviors suggest that there is a risk to the caching animal when burying food in the presence of competitors. Pilfering is assumed to be common, but because an animal who is pilfered from also likely pilfers from others, scatter-hoarding despite the risk of theft is considered a viable and stable strategy (Vander Wall & Jenkins, 2003).

Attempts to quantify the amount of pilfering have mainly assessed the rate of disappearance of human-made caches. In a three-week study of fox squirrels, results suggested pilfering rates of up to 9.4% per day, although a second study used shallower caches, and reported pilfering rates of up to 33% per day (Stapanian & Smith, 1984). Studies of congeneric eastern gray squirrels (*Sciurus carolinensis*) suggested that squirrel-made and human-made caches were removed from the ground at similar rates, although it was not known if the cache owner was also the cache retriever for squirrel-made caches (Thompson & Thompson, 1980). A more recent study of caches made by gray squirrels suggested that all were depleted in less than six days (Steele et al., 2014). However, another study demonstrated that by removing the caching animal from the area immediately after they cached (and thus mimicking predation), caches survived up to 27 days (Steele et al., 2011). This provided evidence that a caching animal holds some advantage in cache recovery but tells us little about what factors led to the pilferage of nuts that were removed in the absence of the animal who original stored them.

Reducing cache density has not shown consistent results in preventing pilferage. In some cases, the loss of human-made caches is reduced by decreasing density (Daly, Jacobs, Wilson, & Behrends, 1992; Male & Smulders, 2008; Male & Smulders, 2007), but in other studies it has had little effect (e.g., Galvez, Kranstauber, Kays, & Jansen, 2009). However, if cache density does increase pilfering, the impact of cache density or of caching close to the caches of other squirrels may be mitigated when pilferers are close relatives. Stapanian and Smith (1978) found that squirrels tended to cache in unique areas, and cached slightly closer to their own previous caches than to those made by other squirrels.

Food theft may be tolerated in animals with overlapping ranges because it is a form of reciprocal exchange that avoids the behavioral costs of cache defense, vigilance, and aggression (Stevens & Stephens, 2002). We currently know very little about the potential effects of kin selection on the pilferage of scatter-hoarded food in free-ranging tree squirrels. One study showed that related male-female and female-female pairs had closer range centers than those of unrelated squirrels. However, the same study found that within a restricted search area (a 50 x 50-m area around the food source), relatedness did not influence the proximity of caches made by different squirrels (Spritzer & Brazeau, 2003). Another study reported a low degree of relatedness within groups of fox squirrels, due to natal dispersal, which is influenced both by age and sex (Koprowski, 1996). Low relatedness would make the question of kin selection less relevant. Population density and dispersal patterns may be adapted to local conditions, however, and it is not clear what group relatedness would be in urban squirrels who are provisioned with food (Penner et al., 2013) or live in fragmented landscapes (Sheperd & Swihart, 1995), both which can impact dispersal.

Reciprocal theft tolerance among related food-storers has been demonstrated in larder-hoarding animals such as woodpeckers (*Melanerpes formicivorus*; Koenig, 1987) and beavers (*Castor canadensis*; Novakowski, 1967). Among scatter-hoarders, there could be fitness benefits in relaxing cache protection strategies in the presence of closely related individuals.

This study had several objectives. The first was to determine if levels of pilfering could be assessed in the field, including identifying specific individual squirrels who pilfer or move caches. If it was possible to observe pilfer events, and determine who was stealing from whom, further study into how behavioral and genetic factors could influence the outcome of caches would be justified.

The second goal was to determine the fate of squirrel-made caches, including how long caches remain where buried, and whether they are pilfered, re-cached, eaten or forgotten. An additional question was whether assessment or cache protection behaviors are related to the outcomes of buried nuts. Despite numerous studies of cache protection, there is little direct evidence that these strategies labeled as cache protection help animals recover their caches, or deter theft by others. We predicted that food assessment and cache protection behaviors should be related to a longer cache life.

The final objective was to examine the population dynamics and heterogeneity of squirrels in the study, including testing the hypothesis that cache proximity and pilferage tolerance could serve as a form of kin selection. Where theft did occur, we predicted there would be an increased likelihood of theft by offspring and other closely related individuals and higher tolerance of pilferage by closely related conspecifics.

## Experiment 1: Testing squirrel responses to stimuli

In order to observe cache movements in the field, we painted 350 caching stimuli (intact hazelnuts) with two coats of yellow non-toxic acrylic paint (Sargent Art, Hazleton, PA). We first tested the squirrels’ ability to discriminate between painted and unpainted hazelnuts to determine whether the paint might make it easier or more difficult for squirrels to locate cached nuts.

### Methods

#### Study Site

The study was conducted outside of Tolman Hall on the University of California, Berkeley campus.

#### Study Animals

Eight free-ranging, marked fox squirrels participated in the study. The research was approved under a protocol submitted to the Animal Care and Use Committee of the University of California, Berkeley.

#### Procedure

Playground sand (Quikrete Cement and Concrete Products, Atlanta, GA) was placed in a 50.8 x 50.8 x 14-cm plastic container at a depth of approximately 5-cm. The container had a latch on one end that allowed the side to be lowered to allow easy access into the box. The apparatus was divided into sixteen 12.7 x 12.7-cm quadrats, numbered from one to sixteen.

Data were collected between October 14 and November 5, 2014. We lured one marked squirrel at a time into the apparatus by calling to them and placing small pieces of peanuts nearby and on top of the sand. Once the squirrel was habituated to entering the apparatus, the peanut pieces were removed.

Four painted nuts, and four unpainted nuts were placed in quadrats chosen by a random number generator (random.com), such that no quadrat had more than one nut in it, and on any given trial, half of the quadrats contained a buried nut. Each hazelnut was covered with enough sand that it could not be detected visually. The focal squirrel was allowed to sniff around and dig in the sand, until it found a hazelnut. Some squirrels did not locate a hazelnut and left.

When a squirrel first located a hazelnut, the following data was recorded: the name of the squirrel, the quadrat the nut was removed from, and whether the nut was painted or unpainted. All squirrels that found hazelnuts carried them away and cached them. Between trials, all nuts were removed from the apparatus, the sand was stirred around to reduce olfactory cues, and nuts were placed in new locations as predetermined by random number generation.

### Results of Experiment 1

Six squirrels completed at least 20 trials. A total of 118 trials were conducted. In 64 (55%) of the trials, the squirrel found a painted hazelnut first; in the remaining 54 trials, the squirrels found an unpainted hazelnut first. Using a binomial probability, this detection rate for painted nuts is not different from chance (binomial test, *p* = .52). From this result, we conclude that the painting of the nuts did not give off odor cues that would influence the difficulty or ease in locating cached nuts when compared to unpainted hazelnuts.

## Experiment 2: Assessing pilferage in the field

The purpose of the pilot study was to determine whether pilferage between individual squirrels could be assessed in the field.

### Methods

#### Study Site

The study was conducted on the University of California, Berkeley campus. This area is relatively open and flat, with oak, pine and other trees, lawns, ivy ground cover and campus buildings. The study area was approximately 0.09 km^2^.

#### Study Animals

Twenty-three free-ranging fox squirrels who regularly frequented the study site participated in the study. All squirrels were individually marked with fur dye (Nyanzol-D, American Color and Chemical Corporation, Charlotte, NC). We chose one adult female (Flame) as the focal subject, because she was frequently seen foraging in the testing area. The research was approved under a protocol submitted to the Animal Care and Use Committee of the University of California, Berkeley.

#### Procedure

The study was conducted between the hours of 10:00 and 16:00 on each weekday from June 16^th^ to July 25^th^, 2014. The caching stimuli were whole hazelnuts, in the shell, which had been painted bright yellow with two coats of non-toxic acrylic paint as in Experiment 1). The focal squirrel recognized the painted hazelnuts as food items, eating or caching all nuts.

On each morning of testing we dispensed up to 15 nuts, one nut at a time, and observed the focal squirrel while she either ate or cached the nut. The number of nuts dispersed was dependent on the presence of the focal squirrel. On some days, she left the study site before all 15 nuts were presented. If a nut was cached, we marked the number of the nut and the location of the cache on a paper map. We also took a GPS waypoint for each cache location. The focal squirrel cached 340 painted hazelnuts.

While nuts were dispersed, researchers noted which other squirrels could be observed in the area. Each day, after dispersing all nuts to the focal squirrel, we used binoculars to observe the squirrels in the study site for several hours each day. The yellow paint allowed for increased visibility of the food items while carried by squirrels. Because the nuts were painted, and all squirrels in the area were marked, when a squirrel was seen moving or eating a yellow hazelnut, we were able to note the identity of the squirrel carrying the painted nut. We also noted where nuts were re-cached.

### Results of Experiment 2

During 125 hours of observation, 102 nuts were observed being moved by a squirrel. We observed the focal squirrel moving and either eating or re-caching 16 of these nuts. The remaining nuts were pilfered by other squirrels, suggesting an overall pilfering rate of at least 25 percent. Our observations suggested that although several individuals were pilfering small amounts from the focal squirrel, some squirrels were more likely to pilfer nuts than others, with two squirrels pilfering 14 and 15 nuts respectively (Figure 1). For 22 caches (25% of stolen caches), nuts were pilfered within 20 minutes of being cached, allowing us to note the specific identity of that cache. Of the two squirrels that frequently stole nuts, one was a juvenile male often spotted in the same tree as the focal squirrel. Behavioral observations suggested this juvenile may have been the offspring of the focal squirrel.

**Figure 1.**
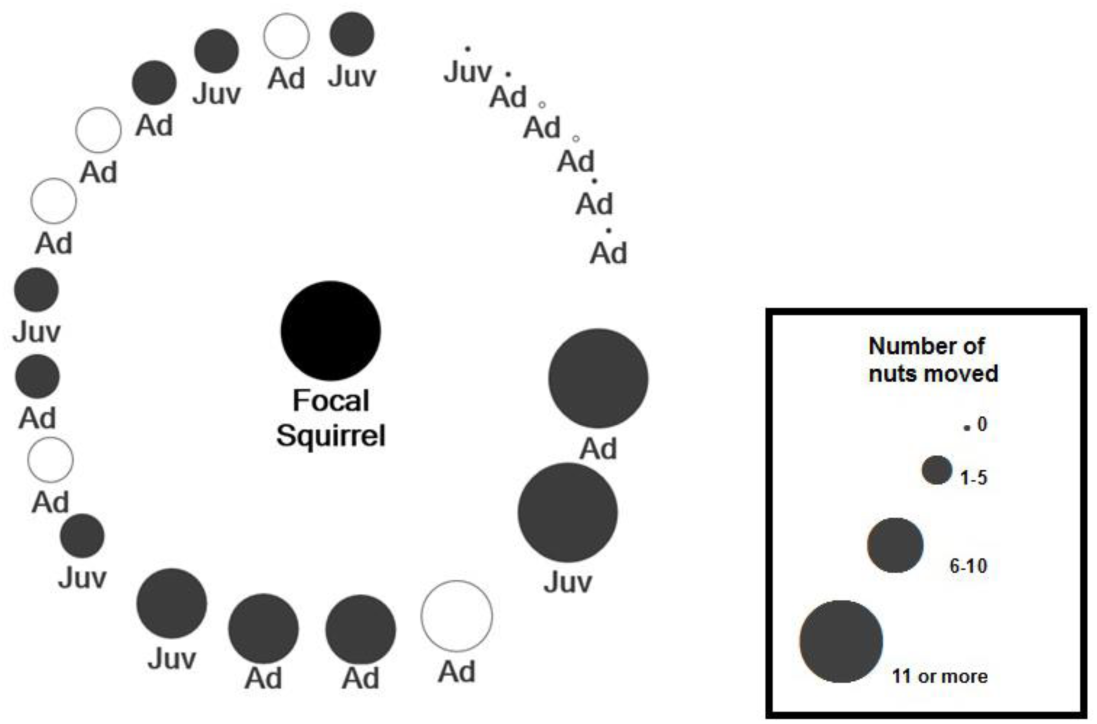
Pilfering of caches made by the focal squirrel. Circles represent theft by either male ● or female ○ adult (Ad) or juvenile (Juv) squirrels. The size of circles represents number of nuts moved.

## Field Study

The pilot data from Experiment 2 demonstrated that it was possible to quantify pilfering in the field, and to identify which squirrels are pilfering specific nuts. The purpose of the current study was to determine (1) what happens over the lifespan of a cache – how many times, where and when is a nut moved before it is finally eaten; (2) the influence of assessment behaviors on cache lifespan and outcomes; and (3) the effect of relatedness of caching behaviors.

### Methods

#### Study Site

The study was conducted on the University of California, Berkeley in the same general area as the previous experiment. The study area was approximately 0.10 km^2^.

#### Study Animals

Nineteen free-ranging fox squirrels who regularly frequented the study site participated in the study. All squirrels were individually marked with Nyanzol-D (American Color and Chemical Corporation, Charlotte, NC). The research was approved under a protocol submitted to the Animal Care and Use Committee of the University of California, Berkeley.

#### Experimental Stimuli

First, 350 hazelnuts were checked to determine that they had no cracks in their shell. A small hole was drilled in each nut using a Dremel Multipro 395 hand-held tool fitted with an 1/16” drill bit. A 12-mm 134.2 kHz pit tag (Biomark, Boise, ID) was placed in each nut, and the hole was filled with Elmer’s wood glue. The surface of the nut was leveled when necessary by ensuring the hole was entirely filled with glue, and scraping away any excess glue. After the glue was dry, the nuts were painted with two coats of bright yellow paint (Sargent Art, Hazleton, PA). Due to experimental oversight, forty of the nuts were painted light green with the same brand of acrylic paint. After the nuts were dried, they were numbered 1 to 20 with a non-toxic marker, and placed in bags of 20 nuts each that were labeled alphabetically, such that each nut had a unique alphanumeric code (for example, A1, A2…B1, B2, etc.). All nuts were scanned with a BioMark HPR Plus reader to verify that their pit tag was functional. We weighed each nut, and entered each nut’s alphanumeric code, pit tag code, and weight into a database.

#### Procedure

A total of 350 pit-tagged nuts were distributed to squirrels from February 11, 2016 until April 5, 2016, between 9:45 and 16:00 hours. On most days, 20 nuts were handed out (10 in the morning and 10 in the afternoon), dependent on weather, lab staffing, and squirrel participation.

A uniquely marked squirrel was solicited for each trial by an experimenter gesturing or calling to the squirrel. One experimenter videotaped all sessions with a Canon FS300 handheld camcorder, noting the squirrel, and alphanumeric code of the nut for each trial. The purpose of videotaping each cache was to record food assessment and cache investment behaviors. If a squirrel could not be easily filmed, experimenters dictated any change in behaviors when they could be observed.

A second experimenter gently tossed the nut on the ground toward the squirrel, and kept records of the subject, time, and the other squirrels present for each trial. The third experimenter scanned the nut at the start and end of the trial with the Biomark HPR Plus reader, which also collected GPS information for the start location and the final cache location.

The squirrel either cached or ate each nut. When the squirrel cached the nut, all experimenters followed the squirrel from a distance until the nut was cached. At that point, the third experimenter scanned the cache location to verify that the nut had been cached and could be detected. The location of the cache was drawn on a map, and the location of the cache was measured from at least two landmarks, noting both distance and bearing (determined by a handheld compass or cell phone compass application) from the landmarks.

Trials were repeated until 10 nuts were handed out for the session or until there were no squirrels available to participate. We alternated between different individual squirrels between trials if multiple subjects were available and willing to participate.

When not handing out nuts, experimenters observed the squirrels to note if there were any cache movements. We used the BioMark HPR Plus to search for previously cached nuts, initially scanning for all cached nuts that had a known location at least every two to three days. Other testing areas were scanned regularly using either the handheld HPR loop antenna, or with a portable antenna that had been mounted on a dolly with wheels to facilitate the rapid search of large, open areas where squirrels often cached. Constraints included weather, staffing, and the battery power of the pit tag reader.

We tracked nuts that had been stolen or re-cached, including their new locations, and if observed, who moved the cache. Cache life was defined as the number of days a cache stayed in its original location. We also recorded any nuts that were detected in a previously unknown location, and then checked those nuts routinely until they disappeared or were still present at the end of the experiment and assumed forgotten. Any new microchips that were detected six months after the end of the experiment were dug up to determine if they were still embedded in a nut or if the nut had been eaten.

All videos of the sessions were coded using The Observer XT (Noldus, Leesburg, VA). There were three video coders, and inter-rater agreement on onset, timing and presence of behaviors between the pairs of coders was high (agreement for coded videos (*n* = 9) averaged Cohen’s kappa, *κ* = .91, range: 0.75 to 1). The variables recorded for all cached nuts included: the number of head flicks for each nut, the amount of time spent paw manipulating, the amount of time spent digging, tamping, and covering the nut, and the amount of time the squirrel spent handling the nut, from initial receipt of the nut until the cache was completed.

One rater assessed the level of concealment of all cache events, whether open (the entire squirrel could be observed caching), partially concealed (more than half of the squirrel was covered by ivy or other plant matter), mostly concealed (less than half of the squirrel’s body could be seen), or completely concealed (none of the squirrel could be observed while caching, such as if the squirrel was caching in a hedge). To determine reliability, a second rater coded 60 of the cache events. Inter-rater agreement for the level of concealment of the cache was *κ* = .84.

GIS data was used to determine the distance traveled for each cache, and the proximity of an individual squirrel’s caches to their own caches and those of other squirrels.

#### Statistical Analyses

All data were analyzed using mixed models in JMP 12.0 (SAS Institute, Cary, NC). All models included squirrel identity as a random effect. The alpha level for all analyses was set at 0.05 and Tukey’s HSD tests were conducted for any pairwise comparisons.

The first model determined if there were effects of nut weight and assessment on distance traveled to cache. A second model examined the effects of assessment and investment behaviors, and social competition on cache life. The independent variables were number of headshakes, time spent paw manipulating, distance traveled, time spent on cache, concealment of cache, time spent digging, tamping, and covering the cache, and the number of other squirrels in the area. A third model was run to determine if squirrels adjusted investment behaviors (digging, tamping, and covering their caches) depending on the level of concealment of the cache location or the presence of other squirrels.

Spatial data were analyzed using ArcGIS version 10.3 (ESRI, Redlands, CA), and JMP Pro12.0 (SAS, Cary, NC. Waypoints were entered into ArcGIS with the WGS 1984 Geographic Coordinate System, and with the State Plane NAD83 California Zone III projection. We calculated the distance traveled for each cache, the proximity of each squirrel’s cache to their own caches and all caches made by other squirrels.

## Experiment 3: DNA Collection and Analysis

In order to assess the effects of relatedness on fox squirrel caching behaviors, hair samples were collected during the same testing period as the rest of the experiment.

### Methods

#### Study Site

The study was conducted on the University of California, Berkeley campus in the same general area as the previous experiments. The study area was approximately .09 km^2^.

#### Study Animals

Hair samples were collected from 14 of the free-ranging, marked fox squirrels who participated in Experiment 3. Hair samples were collected from an additional eight squirrels who were not in the field study. The research was approved under a protocol submitted to the Animal Care and Use Committee of the University of California, Berkeley.

#### Procedure: Hair Collection

Hair collection was based on methods previous described in multiple studies of free-ranging mammals (Finnegan, Hamilton, Perol, & Rochford, 2007; Reiners, Encarnação, & Wolters, 2011). Squirrels were first desensitized to entering a Tomahawk Flush Mount Squirrel trap for food. Both doors of the trap were secured open with zip ties, so the trap would not be set off when an animal entered it. A 60.96 x 20.32-cm black strip of plastic was placed at the bottom of the trap to allow for easy baiting with small pieces of walnuts and peanuts. Once the squirrel entered the trap readily, the trap was set to collect hair.

Experimenters wore latex gloves during all handling of hair collection materials to reduce the risk of contamination. All equipment was sanitized between uses in the field or in the lab with rubbing alcohol. Five 3.58 x 13-cm strips of double-sided Ace brand heavy-duty carpet tape were placed on a piece of PVC tubing (20.32-cm long, diameter 4.11-cm). The tubing was suspended in a storage box by placing it over the center core of a multi-roll tape dispenser (Figure 2a). The storage box and a pair of sanitized tweezers were taken out to the field site.

**Figure 2.**
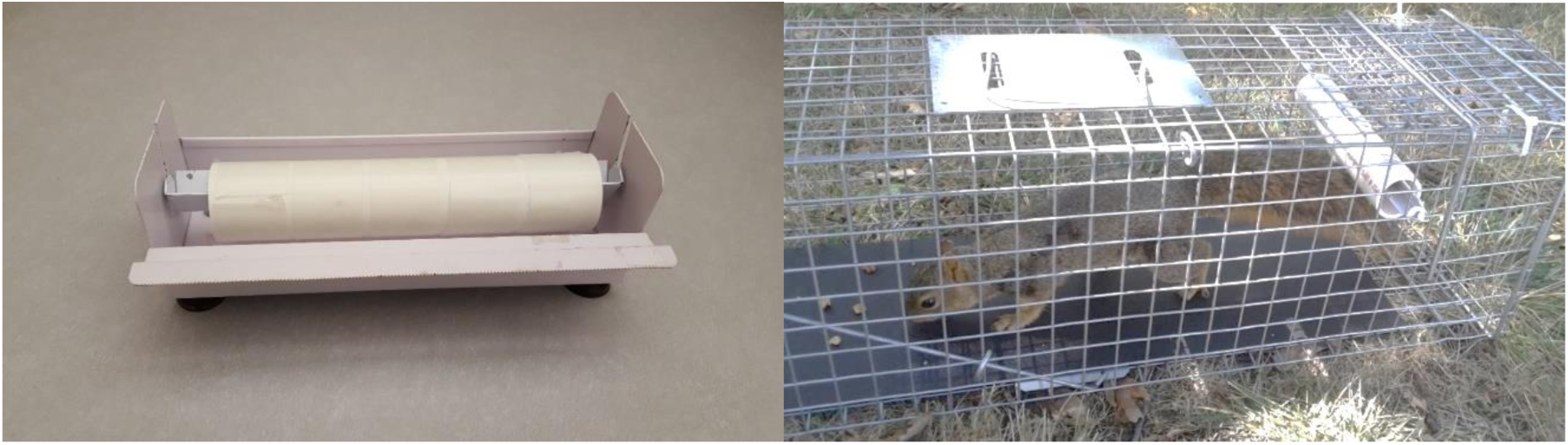
Hair collection procedures. (a) PVC tubing prepared for hair collection. (b) A marked fox squirrel in the trap baited for hair collection.

A marked squirrel was recruited for hair collection. Other squirrels were kept away from the trap by tossing them peanuts. The release liner of the carpet tape was removed with tweezers and PVC tubing was inserted at one end of the trap. The tube was suspended by either a piece of wire affixed to both sides of the trap, or by the core of the tape dispenser (Figure 2b). The tube was suspended low enough that if a squirrel passed underneath it, their tail would touch the exposed tape. The squirrel was lured into the trap several times with walnut pieces, until an adequate number of hairs with follicles were collected from the tail. The tube was removed from the trap and returned to the tape dispenser holder in the plastic storage container. The name and sex of the squirrel, and the date of collection were marked on a label on the container. The container was sealed and stored until hair samples could be processed.

#### Procedure: Preparation of Hair Samples

Hair samples were later prepared for polymer chain reactions (PCR) in a clean environment where no other biological materials were handled. Experimenters wore gloves, a gown, a face mask and a hair net, which were all changed between samples. The surface was sanitized with Sanizide Germicidal Solution (Safetec, Buffalo, NY) and then a large piece of butcher paper was placed on the surface.

The tape dispenser with the hair sample was removed from the plastic storage container. The experimenter removed individual hairs from the tape, inspected them carefully for a follicle, and then cut the hair approximately 2 mm below the follicle. The follicles were placed in an individual Fisherbrand glass threaded 15 x 45-mm, 3.7 mL vial (Fisher Scientific, Chicago, IL) containing ethanol (200 proof Ethyl Alcohol, Spectrum Chemical Mfg. Corp., Gardena, CA). Once an adequate number of hair follicles were collected (generally between 30 and 40 follicles, but fewer if the sample from the squirrel was scant), the tube was sealed and labelled with the squirrel’s name and sex, and the date. In between processing samples, all materials were sanitized with rubbing alcohol, and any other materials (tape, butcher paper, hairs, gloves, gowns, etc.) were disposed of in an individual trash bag that was sealed.

#### DNA Amplification, PCR, and Sequencing

Genetic relatedness and diversity of 15 fox squirrels was inferred from PCR amplification and analysis of 12 microsatellite loci (Table 1). These markers were previously identified as polymorphous in fox squirrels (Fike & Rhodes Jr, 2009). Primers for the 12 loci were acquired from Sigma-Aldrich (The Woodlands, TX).

**Table 1.**
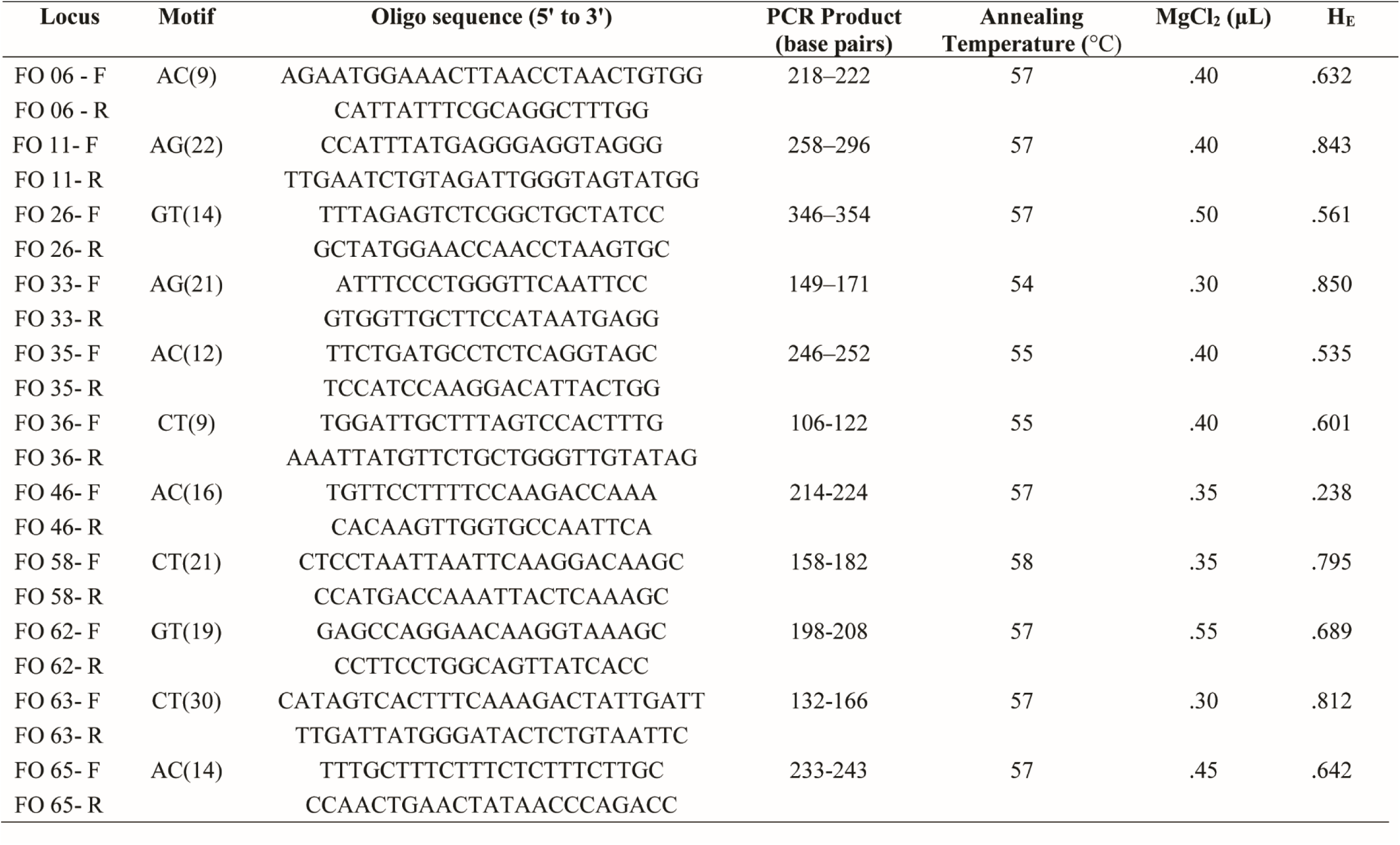
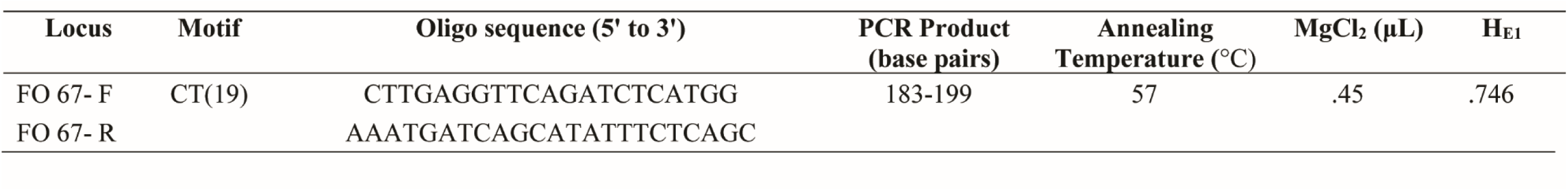
Motifs and oligo sequences for twelve polymorphic microsatellite loci used in the study to determine relatedness among fox squirrels. Annealing temperature, magnesium chloride (MgCl_2_) concentrations used during PCR, expected (H_E_) heterozygosity as reported by Fike and Rhodes (2009).

DNA from 5-10 hair follicles for each individual was extracted using standard methods via a DNEasy Blood and Tissue kit (QIAGEN, Valencia, CA). We amplified the DNA utilizing a polymerase chain reaction process in a BIO-RAD icycler thermal cycler (BIO-RAD, Hercules, CA).

Each 10-μL reaction mixture contained 3 μL of DNA material, 0.3 μL each of the forward and reverse primer, 0.3 - 0.55 μL MgCl_2_ (adjusted for specific primer pairs, see Table 1), 0.25 μL of dNTP, 1.0 μL reaction buffer (Tango, Carlsbad, CA) and 0.12 μL of Taq polymerase (Invitrogen, Carlsbad, CA). The forward primer for primer pairs was fluorescently labeled with either 6-FAM or HEX dye. PCR reactions were run through three steps: (1) denaturation at 95°C for 4 min; (2) 36 cycles of denaturation at 95°C for 45 s, annealing at 54-58°C (adjusted for specific primer pairs, see Table 1) for 30 s and elongation at 72°C for 45 s; and (3) final elongation at 72°C for 10 min.

Successful reactions were prepared for sequencing with 2 μL of PCR product, diluted with 9.8 μL of formamide and combined with 0.2 μL of an internal size standard (LIZ 500, Applied Biosystems, Foster City, CA, U.S.A.). Fragments were determined via sequencing using a Thermo Fisher 3730 DNA Analyzer (Thermo Fisher, Waltham, MA). Base pair lengths were labeled using Geneious 10.1, with the Microsatellite Plugin 1.4 (Biomatters Limited, Newark, NJ).

#### Statistical Analyses

Pairwise relatedness between each pair of subjects in the study were estimated using the program ML-Relate 5.0 (Kalinowski, Wagner, & Taper, 2006), which calculates maximum likelihood estimates of relatedness and the most likely relationship between pairs of individuals. Expected and observed heterozygosity (the probability that an individual will be heterozygous at a given locus) were calculated using the “adegenet” package in R 3.3.0 (Jombart, 2008).

### Results

#### Cache outcomes

A total of 292 nuts were cached. No video was obtained for three caches and some data was missing for these caches. Twenty nuts were eaten at the time they were distributed to squirrels, and 36 nuts had an unknown outcome because the squirrel could not be tracked until they ate or cached the nut.

The average lifespan of a cache was 38.38 days (Median = 4 days, range: 0 to 482 days). The number of nuts cached and cache life by individual are depicted in Table 2. Four hundred and eighty-two days after the start of the experiment, 12 nuts remained in their original cache locations and were assumed forgotten. This suggests an overall forgetting rate of around four percent. An additional 18 nuts remained in new locations that they had been moved to at some point during the experiment, a further loss of six percent. Seven instances of pilfering and one recaching event (by the squirrel “Three”) were observed. Pilfering events between squirrels are noted in Table 4.

**Table 2.** Number of nuts cached by each squirrel, and average cache life (both mean and median) in days.

| Squirrel | Number of nuts cached | Average cache life (days) (SD) | Median cache life |
| --- | --- | --- | --- |
| Biggie<sup>a</sup> | 37 | 55.89 (115.75) | 4 |
| Billy Ray | 3 | 18.00 (26.00) | 4 |
| Blake | 18 | 28.89 (39.40) | 11.5 |
| Chubs<sup>b</sup> | 23 | 91.65 (177.05) | 5 |
| Curly | 1 | 24.00 (NA) | 24 |
| December | 1 | 0.00(NA) | 0 |
| Fermata | 7 | 23.57 (24.41) | 22 |
| Flame | 16 | 26.31 (69.75) | 4 |
| Gwen | 1 | 3 (NA) | 3 |
| Jewel | 4 | 2.75 (3.50) | 1 |
| Joker | 5 | 2.40 (1.14) | 2 |
| Mermaid | 1 | 1.00 (NA) | 1 |
| Roger | 21 | 21.29 (61.26) | 5 |
| Scarf | 16 | 29.25 (63.69) | 4 |
| Squiggle<sup>c</sup> | 43 | 41.40 (105.11) | 4 |
| Stool<sup>a</sup> | 28 | 67.54 (136.01) | 9.5 |
| Stovetop | 15 | 4.53 (5.83) | 3 |
| Teddy Bear | 2 | 2.00 (1.41) | 2 |
| Three | 47 | 24.57 (73.52) | 3 |
| Walter | 3 | 2.33 (2.08) | 3 |

<sup>a</sup>Forgot three caches
<sup>b</sup>Forgot four caches
<sup>c</sup>Forgot two caches

**Table 3.** Expected (H_E_) and observed (H_O_) heterozygosities at the twelve loci analyzed.

| Locus | $H_E$ | $H_O$ |
| --- | --- | --- |
| F06 | 0.64 | 1.00 |
| F26 | 0.71 | 0.95 |
| F11 | 0.92 | 1.00 |
| F33 | 0.73 | 1.00 |
| F35 | 0.71 | 0.59 |
| F36 | 0.60 | 0.71 |
| F46 | 0.71 | 0.95 |
| F58 | 0.81 | 1.00 |
| F62 | 0.34 | 0.20 |
| F63 | 0.68 | 0.90 |
| F65 | 0.64 | 0.86 |
| F67 | 0.72 | 0.95 |

**Table 4.** Probabilities of relatedness between individuals in the study, as calculated by ML-Relate.

|  | Biggie | Roger | Teddy | Fermata | Walter | Stool | Joker | Blake | Flame | Jewel | Three | Squiggle | Curly | Chubs |
| --- | --- | --- | --- | --- | --- | --- | --- | --- | --- | --- | --- | --- | --- | --- |
| Biggie | 1 |  |  |  |  |  |  |  |  |  |  |  |  |  |
| Roger | 0.07 | 1 |  |  |  |  |  |  |  |  |  |  |  |  |
| Teddy | 0.08 | 0.13<sup>c*</sup> | 1 |  |  |  |  |  |  |  |  |  |  |  |
| Fermata | 0 | 0.06 | 0.36<sup>b</sup> | 1 |  |  |  |  |  |  |  |  |  |  |
| Walter | 0.19 | 0.03 | 0.11 | 0.13 | 1 |  |  |  |  |  |  |  |  |  |
| Stool | 0.19 | 0.08 | 0.14 | 0 | 0.25 | 1 |  |  |  |  |  |  |  |  |
| Joker | 0.01 | 0 | 0 | 0.08 | 0.2 | 0.13 | 1 |  |  |  |  |  |  |  |
| Blake | 0.27 | 0 | 0 | 0.16 | 0 | 0 | 0.15 | 1 |  |  |  |  |  |  |
| Flame | 0.51<sup>b</sup> | 0 | 0 | 0 | 0.02 | 0.05 | 0 | 0.70<sup>b</sup> | 1 |  |  |  |  |  |
| Jewel | 0 | 0 | 0.09 | 0.31 | 0 | 0 | 0 | 0.56<sup>a</sup> | 0.43<sup>a</sup> | 1 |  |  |  |  |
| Three | 0.29<sup>b*</sup> | 0.30<sup>c</sup> | 0 | 0 | 0.16 | 0.38<sup>a</sup> | 0 | 0 | 0.12 | 0 | 1 |  |  |  |
| Squiggle | 0.28 | 0 | 0.10<sup>*</sup> | 0.12 | 0.24 | 0.37<sup>b</sup> | 0 | 0.08 | 0.24 | 0.16<sup>*</sup> | 0 | 1 |  |  |
| Curly | 0 | 0 | 0 | 0.09 | 0 | 0 | 0 | 0.08 | 0.18 | 0.04 | 0 | 0.21<sup>c</sup> | 1 |  |
| Chubs | 0.02<sup>*</sup> | 0 | 0 | 0 | 0 | 0.22<sup>b</sup> | 0.24<sup>c</sup> | 0.4 | 0.13 | 0.03 | 0 | 0.14<sup>c</sup> | 0 | 1 |

<sup>a</sup>Likely parent-offspring relationship
<sup>b</sup>Likely full-sibling relationship
<sup>c</sup>Likely half-sibling relationship
<sup>\*</sup>Pilfering event observed between these two individuals

The only variable that was related to the length of time a cache stayed in its original location was the level of concealment (*F*(3, 232) = 3.32, *p* = .021) such that caches that were placed in mostly concealed areas had longer cache lives (*n* = 39, *M* = 93.38 days, 95% CI [40.92, 145.83 days], Median = 8 days) than caches placed in open (*n* = 175, *M* = 32.80 days, 95% CI [19.81, 45.77 days], Median = 4 days) or partially concealed areas (*n* = 56, *M* = 21.30 days, 95% CI [5.33, 37.27 days], Median 3.5 days). Caches placed in totally concealed areas had a lifespan of 26 days (*n* = 20, 95% CI [-4.47, 56.47 days], Median = 5.5 days) and were not statistically different from other cache concealment categories.

Weight and the number of headshakes were weakly related to the distance from the food source that a squirrel traveled to cache, such that heavier nuts and more headshakes were associated with a longer distance traveled but the effect in both cases was not statistically significant (weight: *F*(1, 275) = 3.14, *p* = .08; headshakes *F*(1, 75.78) = 2.91, *p* = .09).

Finally, squirrels adjusted cache protection behaviors depending on the level of conspicuousness of the cache. They spent more time caching nuts when in open locations (*F*(3, 269.4) = 3.76, *p* = .011), or when other squirrels were present (*F*(7, 265.2) = 2.72, *p* = .010; Figure 3). Squirrels spent the most time digging (*F*(3, 254.1) = 4.43, *p* = .005), and covering their caches (*F*(3, 256.5) = 13.68, *p* < .001) when they cached in an open location, and spent the least amount of time on all cache protection behaviors (digging, tamping, and covering caches) when in a concealed location. See Figure 4.

**Figure 3.**
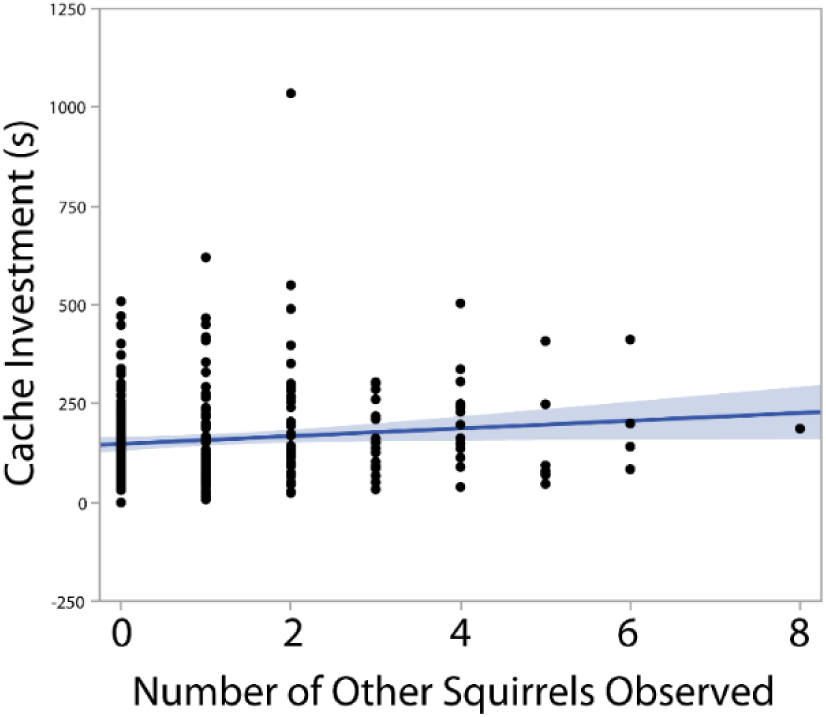
Total cache time (seconds) in the presence of other squirrels. Squirrels tend to spend more time caching as the number of competitors (other squirrels) increases.

**Figure 4.**
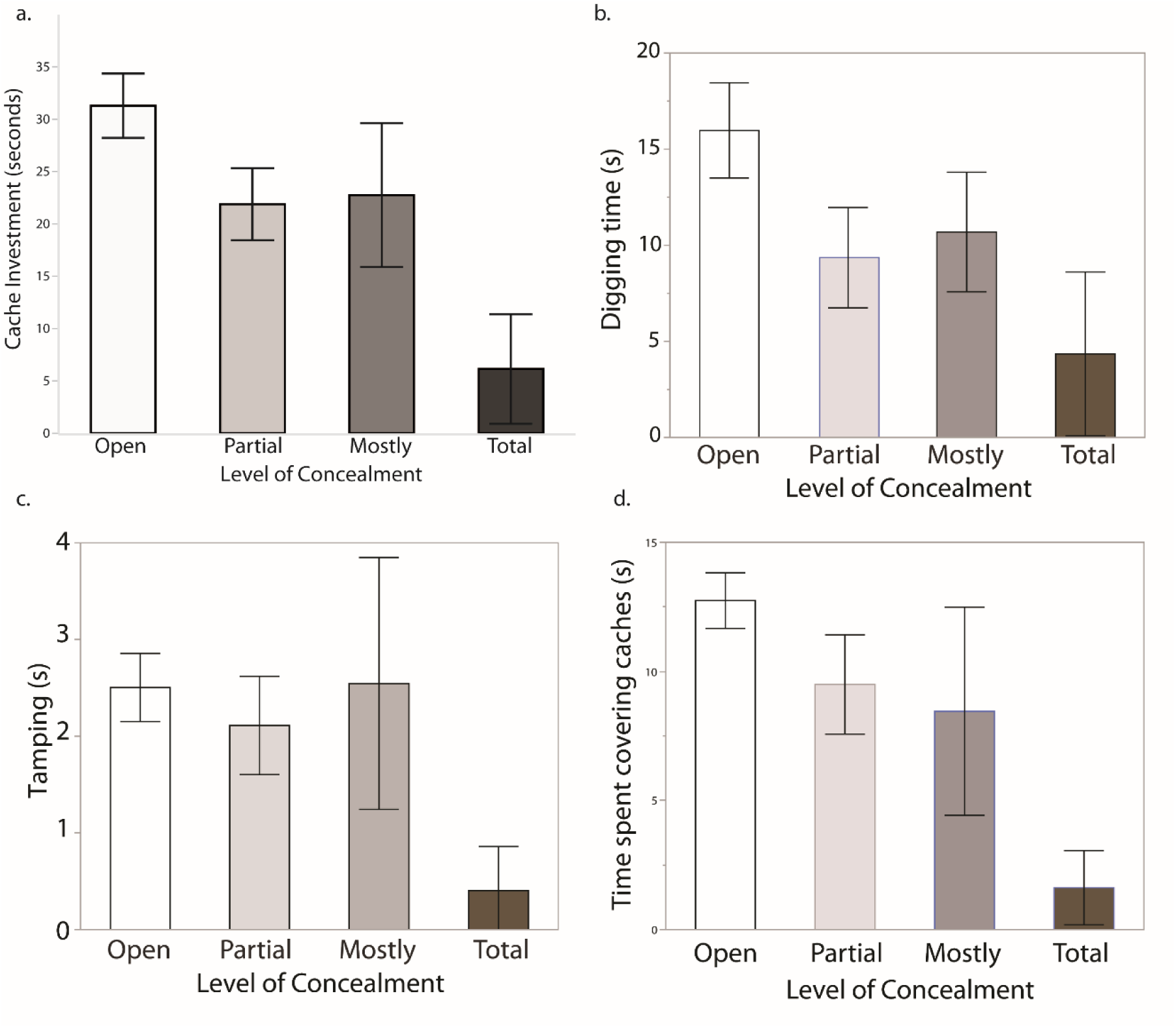
Cache investment and protection at different levels of cache conspicuousness. Squirrels spent more total time caching (a), more time digging (b), and more time covering (d) caches made in open locations compared to completely concealed locations. Squirrels spent the least amount of time tamping caches (c) made in completely concealed locations.

#### Microsatellite analysis

The number of alleles per locus ranged from 3 to 16, and single locus heterozygosities ranged from 0.20 to 0.92 (Table 3), suggesting an overall high level of genetic diversity in the tested population. From 10000 randomized simulations performed in ML-Relate, a possible heterozygote deficiency was found at one loci (F62, *p* = .059; Table 3). Observed heterozygosity was slightly higher than expected (*t*_11_ = −2.09, *p* = .06).

Based on estimates of the most likely relationships between individuals (unrelated, half siblings, full siblings or parent-offspring), there were likely six full siblings, five half siblings, and three parent-offspring relationships between the fourteen individuals in the study for whom we had DNA samples (see Table 4).

#### Spatial distribution of caches

Geospatial data was used to assess the proximity of a squirrel’s caches to their own caches, and to those of other squirrels, based on relatedness between individuals. When treated as a continuous variable, there was an negative linear relationship between probability of relatedness and cache distance (*F*(1, 105) = 9.77, *p* = .002, Figure 5a), but this effect was largely driven by the inclusion of the distance each squirrel tended to cache from their own other caches.

**Figure 5.**
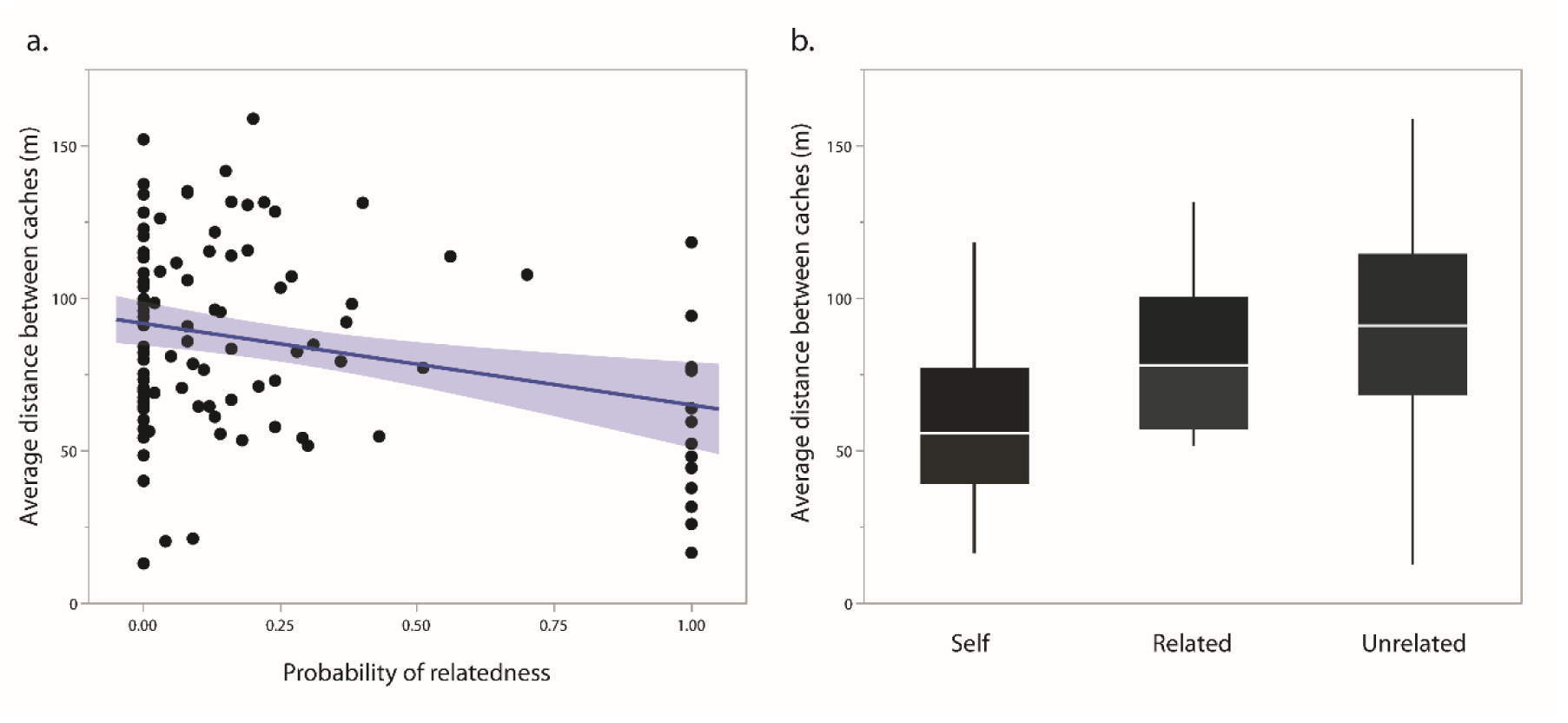
The relationship between relatedness and distance between caches. Relatedness decreases distance between caches (a); squirrels tend to cache closer to their own previously made caches than to those of other squirrels.

When assessed as a categorical variable (self, related, unrelated), there were differences between groups on average distance between caches (*F*(2, 99.42) = 10.71, *p* < .001). Squirrels tended to cache closer to their own caches (*M* = 59.14 m, 95% CI [44.87, 73.41]) than to those of other squirrels, particularly when compared to those of unrelated squirrels (*M* = 91.28 m, 95% CI [84.26, 98.3]). The average distance between related squirrels was *M* = 81.93 m, 95% CI [67.37, 96.49]. See Figure 5b.

Squirrels also tended to disperse their caches more as the experiment continued. The distance traveled from food source to cache increased during each consecutive week of the experiment, (*F*(1, 290) = 7.70, *p* = .006, Figure 6). The density of nuts decreased as the experiment continued (Table 5), although squirrels continued to cache in the central area that they cached in during week 1 throughout the remainder of the experiment (Figure 7).

**Figure 6.**
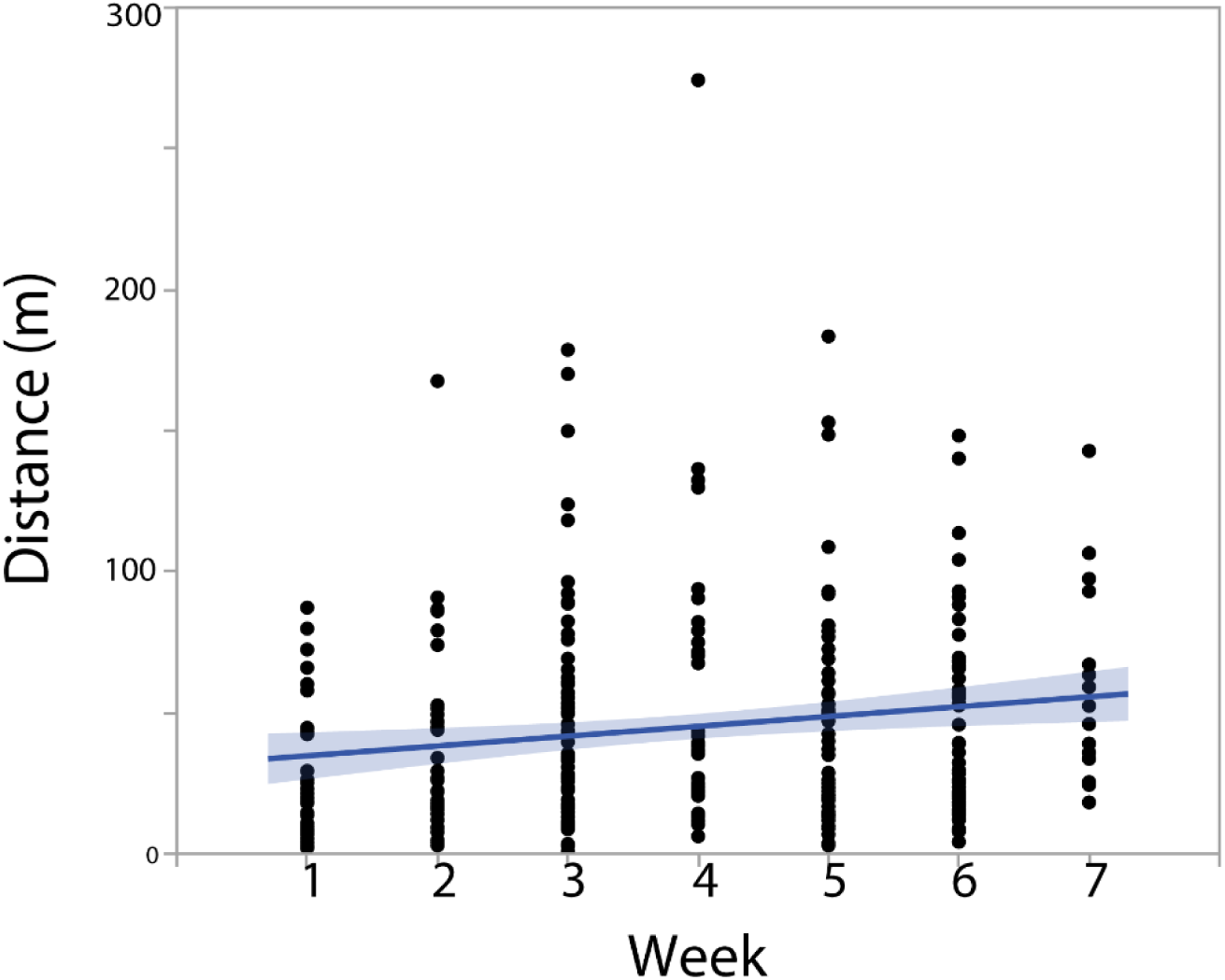
Distance traveled for each cache buried by each week of the experiment. Squirrels increased distance traveled from the food source as the experiment continued.

**Figure 7.**
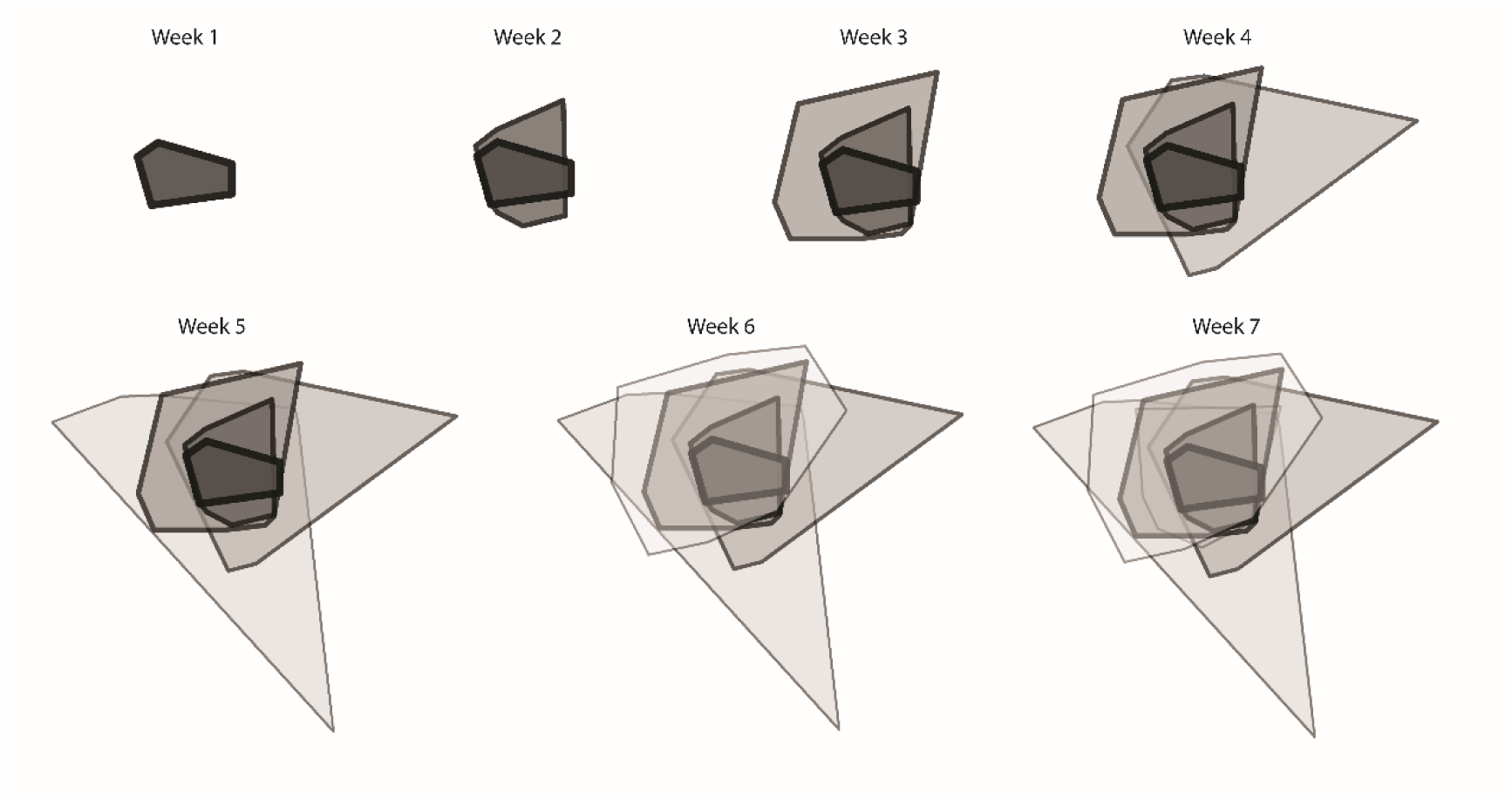
Polygons depicting the minimum bounding geometry for caches made by all squirrels for each week of the experiment. Squirrels utilized a larger area to cache in as the experiment continued, but also continued to cache in a core central area.

**Table 5.** Nearest Neighbor Distances throughout the experiment. NN ratios larger than one indicate nuts that were cached at a lower density than expected if randomly distributed. Observed distances between nuts tended to increase as the experiment continued.

| Week | NN Ratio | Z-statistic | p-value | Observed distance (m) | Expected distance (m) |
| --- | --- | --- | --- | --- | --- |
| 1 | 1.04 | 0.42 | .680 | 8.43 | 8.13 |
| 2 | .88 | -1.34 | .180 | 9.41 | 10.64 |
| 3 | .88 | -1.89 | .060 | 10.48 | 11.87 |
| 4 | .84 | -1.71 | .090 | 20.44 | 24.28 |
| 5 | .80 | -2.75 | .006 | 19.37 | 24.20 |
| 6 | 2.03 | 1.15 | .040 | 21.34 | 18.50 |
| 7 | 1.52 | 4.08 | <.001 | 32.45 | 21.38 |

## Discussion

The results of this study suggest that the most important factor contributing to the fate of caches made by fox squirrels, strictly measured as how long a cache remained in its original location, is the conspicuousness of the cache. Caches that were placed in open areas were moved sooner than other caches. Squirrels also spent more time digging, tamping and covering caches in open areas, compared to more concealed caches.

This study also supported previous findings that squirrels are sensitive to food item value and the social context when caching. Squirrels showed a tendency to travel further for heavier hazelnuts, even though the range of nut weights in this study was very small (*x* = 3.94 g, range: 2.3 to 5.5 g). Several studies that have shown that tree squirrels tend to travel further for heavier nuts, nuts that provide more nutritional content, and nuts that are at lower risk of perishability (Delgado et al., 2014; Moore et al., 2007; Preston & Jacobs, 2009; Stapanian & Smith, 1984; Steele, Hadj-Chikh, Agosta, Smallwood, & Tomlinson, 1996), and this study demonstrates that this may also happen on a very fine-grained scale, even when there are small differences in quality between food items.

Squirrels traveled further away from the food source to cache when greater numbers of competitors were present. They also showed some tendencies to cache closer to their own previously made caches, and closer to the caches made by related squirrels than unrelated squirrels. This supports that squirrels, although generally considered solitary (Steele & Koprowski, 2001) are sensitive to the social context they are caching within.

It has been assumed that the time squirrels spend covering caches is somehow related to preventing conspecific theft. Covering caches has been previously described as a method of disguising caches or as cache protection (e.g., Delgado et al., 2014; Hopewell & Leaver, 2008; Steele et al., 2008). The current study showed that more time covering caches was not a predictor of cache life and in fact the inverse may be true. Squirrels spent more time covering caches that were in open areas, and those caches also tended to stay in place the shortest amount of time. In order to fully understand this effect, it would be necessary to assess the effect of substrate on covering time; it is possible that caches in open areas were placed in a more compact, tighter substrate that required more digging and covering than a looser soil.

If in fact caches are recovered by the squirrel who cached them, then cache covering may serve as protection. But even if the food-storing animal retrieves their own caches, the function of covering needs to be disentangled between different possible hypotheses. Covering caches could provide protection by creating scent cues or consolidating the memory of the food-storer, making retrieval easier for the caching animal. It could also provide protection by making it more difficult for a competitor to find and pilfer a cache.

However, in Experiment 1, 25% of pilfered nuts were stolen shortly after they were cached. This suggests that squirrels may be observing each other cache; in which case, spending more time covering could provide a signal to competitors that a nut is being buried, and give pilferers more time to observe the cache location. The function of cache covering behavior merits further exploration, but most importantly how the outcome of caches is related to covering behavior needs to be determined.

The results of this study demonstrated that pilfering between individual squirrels can be quantified in the field. Unfortunately, we were unable to observe many instances of pilfering or recaching in the final experiment. Given the results from the pilot study, this was surprising. However, in the pilot study, we only provided one squirrel with nuts to cache. This limited the area that needed to be observed, as the focal squirrel cached most of the nuts she was provided with in a central area. Provisioning her with nuts each day may have artificially inflated the pilfering rate by changing the caching behavior of only one individual in the study area.

Conversely, in the final study, because several squirrels were caching, the cache areas were distributed across a larger area of the testing area (Figure 7), which made observation difficult as the experiment continued. Furthermore, because we were providing squirrels with nuts in both the morning and afternoon, this limited our total observation time. Because many nuts were moved within a short period of time, the lack of pilfer and recache observations does not suggest that squirrels were not pilfering and recaching nuts; they just did so in times and places that were not being directly observed.

A previous study suggested that the experimental provision of food for squirrels could increase pilferage (Penner et al., 2013). In that study, researchers first provided squirrels with ad libitum food in one plot and did not offer food in a control plot. Later, pecans were buried at identical densities in both plots, and pilfering was statistically higher in the previously provisioned plot. We have not quantified how providing the squirrels with food in our study may have inflated pilferage; however, the current study did not include any provision of food prior to the experiment. During the study, squirrels were provided with nuts primarily where they were observed, thus the provisioning location frequently changed. No specific area of the study site should have been seen as more desirable for foraging or searching for previously made caches.

Squirrels buried the majority (almost 60%) of their caches in an open area, which suggests there may be some benefits to caching in an open area, such as ease of retrieval for short term storage. That said, five out of seven of the observed pilferage events were of nuts were originally cached in open areas. In a previous study (Steele et al., 2014), human-made caches under canopy were moved more than caches made in the open. Based on the limited data we acquired in this study, fox squirrel caches in open areas may be pilfered more frequently. It is possible that since gray squirrels spend more time under canopy in comparison to fox squirrels (Steele & Koprowski, 2001), they were more likely to discover human-made caches under canopy than in the open.

In the current study, half of all cached nuts were moved within four days of being buried. That said, 25% of cached nuts had a life span longer than 20 days. A previous study of squirrel-cached acorns found that of 57 cached nuts, all were moved between one and six days after burial. No relationship was found between cache life and distance nuts were buried from cover. Because it is unknown in both studies if short lifespans are due to pilfering or recaching, it is difficult to say whether this life span is beneficial or detrimental to caching animals.

Approximately 10% of cached nuts remained in place a year after they were cached or re-cached. Based on observations of nuts that were dug up six months after the end of the experiment, they were likely no longer edible. Perhaps the squirrels could detect this and abandoned caches, or these forgotten caches may represent what percent of nuts is typically forgotten by food-storers. Cahalane (1942) found that fewer than two percent of nuts buried by fox squirrels were forgotten over the winter, but as he marked caches with stakes, he may have provided additional visual cues to the original food-storers or to pilferers that made these nuts easier to locate.

A key function of seed dispersers is to propagate tree species (Price & Jenkins, 1986; Sun & Zhang, 2013; Vander Wall, 1990), and squirrels have co-evolved with their food sources (Stapanian & Smith, 1978; Steele, Wauters, Larsen, & Forget, 2004; Vander Wall, 2010). Thus, some forgetting of cached nuts provides benefits to both the tree species, and the food-storer, in terms of guaranteeing future food sources for kin. It is not possible to test the duration of memory for caches with human-made caches, and so pit-tagging of nuts provides an excellent methodology for further testing what percent of nuts may be forgotten by caching animals.

The microsatellite analysis of DNA collected for subjects in this study demonstrated that despite a fragmented habitat, human-made structures, and likely artificial supplementation of food, there is a similar level of genetic diversity among the study population as the populations of fox squirrels sampled in their native habitat (Fike & Rhodes Jr, 2009). We were able to use a non-invasive method to obtain hair samples from free-ranging squirrels that provided adequate DNA for sequencing and analysis. This analysis found expected levels of heterozygosity at 11 out of 12 loci.

More importantly, microsatellite analysis allowed us to explore how relatedness impacts caching behavior. Although we were not able to determine the relationship between probability of relatedness and likelihood of pilfering between individuals, results suggested that squirrels may cache nuts closer to caches made by relatives than unrelated squirrels. If squirrels are more likely to pilfer within or close to their caching territory, then this would suggest some form of kin selection could be at work. This could also prevent pilfering from non-related individuals. Given the small sample size, and the fact that the effect was small, we should interpret these results with some caution; further studies should examine this possibility in much more detail.

Ideally, this study would be replicated with fewer caching subjects and more time to observe individual cache movements. Alternatively, the focal squirrel could be rotated, testing just one individual at a time, to allow for a more fine-grained observations and analysis of the relationship between caching behaviors, relatedness and cache fate. Ideally, hair samples would be collected from all participating squirrels in the study, in addition to sampling squirrels in other locations surrounding the test area, to better assess the level of dispersal among this population of squirrels.

To summarize, this study established or validated several methods for testing the caching behavior and population dynamics of a group of free-ranging, scatter-hoarding tree squirrels. The results demonstrate the flexibility of squirrels when storing food and show that they adjust behaviors according to several environmental and social factors. They also point to the need for a greater understanding of how these behaviors are related to the outcomes of caches that are stored for future use, a question that turned out to be much more challenging to answer than anticipated.

## Acknowledgements

This work was funded by the National Science Foundation Graduate Research Fellowship, and a National Science Foundation Doctoral Dissertation Improvement Grant. We would like to acknowledge Aslan Brown, Daniel Petrie, Peter Buto, Simon Campo, Aryan Sharif, Samuel Kim, Vanessa Alschuler, Lisa Yoen Lee, Alan Xu, Stephanie Kuo, Sylvia Chen, Emani Harris, Marisa Fong, Amy Hseuh, Paul Kim, Breana Martinez, Sarina Utamsing, Kristin Witte, Nicole Breer, Jihoon Park, Luke Strgar, Alyssa Alvarez, Aaron Teixeira, and Tiffany Chan for helping with data collection.

## References

1. Cahalane, V. H. (1942). Caching and recovery of food by the western fox squirrel. The Journal of Wildlife Management, 6, 338–352.

2. Dally, J. M., Clayton, N. S., & Emery, N. J. (2006). The behaviour and evolution of cache protection and pilferage. Animal Behaviour, 72, 13–23. doi:10.1016/j.anbehav.2005.08.020

3. Dally, J. M., Emery, N. J., & Clayton, N. S. (2004). Cache protection strategies by western scrub-jays (*Aphelocoma californica*): Hiding food in the shade. Proceedings: Biological Sciences, 271, S387–S390. doi:10.1098/rsbl.2004.0190

4. Dally, J. M., Emery, N. J., & Clayton, N. S. (2005). Cache protection strategies by western scrub-jays, *Aphelocoma californica*: Implications for social cognition. Animal Behaviour, 70, 1251–1263.

5. Daly, M., Jacobs, L. F., Wilson, M. I., & Behrends, P. R. (1992). Scatter hoarding by kangaroo rats (*Dipodomys merriami*) and pilferage from their caches. Behavioral Ecology, 3, 102–111. doi:10.1093/beheco/3.2.102

6. Delgado, M. M., Nicholas, M., Petrie, D. J., & Jacobs, L. F. (2014). Fox squirrels match food assessment and cache effort to value and scarcity. PLoS ONE, 9, e92892. doi:10.1371/journal.pone.0092892

7. Emery, N., Dally, J., & Clayton, N. (2004). Western scrub-jays (*Aphelocoma californica*) use cognitive strategies to protect their caches from thieving conspecifics. Animal Cognition, 7, 37–43. doi:10.1007/s10071-003-0178-7

8. Fike, J. A., & Rhodes Jr, O. E. (2009). Characterization of twenty-six polymorphic microsatellite markers for the fox squirrel (*Sciurus niger*) and their utility in gray squirrels (*Sciurus carolinensis*) and red squirrels (*Tamiasciurus hudsonicus*). Conservation Genetics, 10, 1545–1548.

9. Finnegan, L., Hamilton, G., Perol, J., & Rochford, J. (2007). The use of hair tubes as an indirect method for monitoring red and grey squirrel populations. Paper presented at the Biology and Environment: Proceedings of the Royal Irish Academy.

10. Galvez, D., Kranstauber, B., Kays, R. W., & Jansen, P. A. (2009). Scatter hoarding by the Central American agouti: A test of optimal cache spacing theory. Animal Behaviour, 78, 1327–1333.

11. Goodwin, D. (1956). Further observations on the behaviour of the jay *Garrulus glandarius*. Ibis, 98, 186–219.

12. Hopewell, L. J., & Leaver, L. A. (2008). Evidence of social influences on cache-making by grey squirrels (*Sciurus carolinensis*). Ethology, 114, 1061–1068. doi:10.1111/j.1439-0310.2008.01554.x

13. Hopewell, L. J., Leaver, L. A., & Lea, S. E. G. (2008). Effects of competition and food availability on travel time in scatter-hoarding gray squirrels (*Sciurus carolinensis*). Behavioral Ecology, 19, 1143–1149.

14. Jablonski, P. G., Fuszara, E., Fuszara, M., Jeong, C., & Lee, W. Y. (2015). Proximate mechanisms of detecting nut properties in a wild population of Mexican Jays (*Aphelocoma ultramarina*). Journal of Ornithology, 156, 163–172.

15. James, P. C., & Verbeek, N. A. (1984). Temporal and energetic aspects of food storage in northwestern crows. Ardea, 72, 207–215.

16. Jokinen, S., & Suhonen, J. (1995). Food caching by willow and crested tits: A test of scatterhoarding models. Ecology, 76, 892–898.

17. Jombart, T. (2008). adegenet: a R package for the multivariate analysis of genetic markers. Bioinformatics, 24, 1403–1405.

18. Kalinowski, S. T., Wagner, A. P., & Taper, M. L. (2006). ML-Relate: A computer program for maximum likelihood estimation of relatedness and relationship. Molecular Ecology Notes, 6, 576–579.

19. Kislalioglu, M., & Gibson, R. N. (1976). Prey ‘handling time’ and its importance in food selection by the 15-spined stickleback, *Spinachia spinachia*. Journal of Experimental Marine Biology and Ecology, 25, 151–158. doi:10.1016/0022-0981(76)90016-2

20. Koenig, W. D. (1987). Population ecology of the cooperatively breeding acorn woodpecker. Princeton, NJ: Princeton University Press.

21. Koprowski, J. L. (1996). Natal philopatry, communal nesting, and kinship in fox squirrels and gray squirrels. Journal of Mammalogy, 77, 1006–1016.

22. Lahti, K., & Rytkönen, S. (1996). Presence of conspecifics, time of day and age affect willow tit food hoarding. Animal Behaviour, 52, 631–636.

23. Langen, T. A., & Gibson, R. M. (1998). Sampling and information acquisition by western scrub-jays, *Aphelocoma californica*. Animal Behaviour, 55, 1245–1254. doi:10.1016/S0003-3472(98)90000-8

24. Leaver, L., Hopewell, L., Caldwell, C., & Mallarky, L. (2007). Audience effects on food caching in grey squirrels (*Sciurus carolinensis*): Evidence for pilferage avoidance strategies. Animal Cognition, 10, 23–27. doi:10.1007/s10071-006-0026-7

25. Ligon, J. D., & Martin, D. J. (1974). Piñon seed assessment by the piñon jay, *Gymnorhinus cyanocephalus*. Animal Behaviour, 22, 421–429. doi:10.1016/S0003-3472(74)80040-0

26. Male, L., & Smulders, T. (2008). Hyper-dispersed cache distributions reduce pilferage: A laboratory study. Journal of Avian Biology, 39, 170–177.

27. Male, L. H., & Smulders, T. V. (2007). Hyperdispersed cache distributions reduce pilferage: A field study. Animal Behaviour, 73, 717–726.

28. Melin, A., Fedigan, L., Hiramatsu, C., Hiwatashi, T., Parr, N., & Kawamura, S. (2009). Fig foraging by dichromatic and trichromatic *Cebus capucinus* in a tropical dry forest. International Journal of Primatology, 30, 753–775. doi:10.1007/s10764-009-9383-9

29. Moore, J. E., McEuen, A. B., Swihart, R. K., Contreras, T. A., & Steele, M. A. (2007). Determinants of seed removal distance by scatter-hoarding rodents in deciduous forests. Ecology, 88, 2529–2540. doi:10.1890/07-0247.1

30. Novakowski, N. S. (1967). The winter bioenergetics of a beaver population in northern latitudes. Canadian Journal of Zoology, 45, 1107–1118.

31. Penner, J. L., Zalocusky, K., Holifield, L., Abernathy, J., McGuff, B., Schichtl, S., … Moran, M. D. (2013). Are high pilferage rates influenced by experimental design? The effects of food provisioning on foraging behavior. Southeastern Naturalist, 12, 589–598.

32. Preston, S. D., & Jacobs, L. F. (2009). Mechanisms of cache decision making in fox squirrels (*Sciurus niger*). Journal of Mammalogy, 90, 787–795. doi:10.1644/08-mamm-a-254.1

33. Price, M. V., & Jenkins, S. H. (1986). Rodents as seed consumers and dispersers. In D. R. Murray (Ed.), Seed Dispersal. New York, NY: Academic Press.

34. Reiners, T. E., Encarnação, J. A., & Wolters, V. (2011). An optimized hair trap for non-invasive genetic studies of small cryptic mammals. European Journal of Wildlife Research, 57, 991–995.

35. Shaw, R., & Clayton, N. (2014). Pilfering Eurasian jays use visual and acoustic information to locate caches. Animal Cognition, 17, 1281–1288. doi:10.1007/s10071-014-0763-y

36. Shaw, R. C., & Clayton, N. S. (2013). Careful cachers and prying pilferers: Eurasian jays (*Garrulus glandarius*) limit auditory information available to competitors. Proceedings of the Royal Society B: Biological Sciences, 280, 20122238. doi:10.1098/rspb.2012.2238

37. Sheperd, B. F., & Swihart, R. K. (1995). Spatial dynamics of fox squirrels (*Sciurus niger*) in fragmented landscapes. Canadian Journal of Zoology, 73, 2098–2105. doi:10.1139/z95-247

38. Spritzer, M. D., & Brazeau, D. (2003). Direct vs. indirect benefits of caching by gray squirrels (*Sciurus carolinensis*). Ethology, 109, 559–575. doi:10.1046/j.1439-0310.2003.00897.x

39. Stapanian, M. A., & Smith, C. C. (1978). A model for seed scatterhoarding: Coevolution of fox squirrels and black walnuts. Ecology, 59, 884–896.

40. Stapanian, M. A., & Smith, C. C. (1984). Density-dependent survival of scatterhoarded nuts: An experimental approach. Ecology, 65, 1387–1396.

41. Steele, M., Wauters, L., Larsen, K., & Forget, P. (2004). Selection, predation and dispersal of seeds by tree squirrels in temperate and boreal forests: Are tree squirrels keystone granivores? In J. E. Lambert, P. E. Hulme, & S. B. Vander Wall (Eds.), Seed fate: Predation, dispersal, and seedling establishment (pp. 205–221). Cambridge, MA: CABI Publishing.

42. Steele, M. A., Bugdal, M., Yuan, A., Bartlow, A., Buzalewski, J., Lichti, N., & Swihart, R. (2011). Cache placement, pilfering, and a recovery advantage in a seed-dispersing rodent: Could predation of scatter hoarders contribute to seedling establishment? Acta Oecologica, 37, 554–560. doi:10.1016/j.actao.2011.05.002

43. Steele, M. A., Contreras, T. A., Hadj-Chikh, L. Z., Agosta, S. J., Smallwood, P. D., & Tomlinson, C. N. (2014). Do scatter hoarders trade off increased predation risks for lower rates of cache pilferage? Behavioral Ecology, 25, 206–215. doi:10.1093/beheco/art107

44. Steele, M. A., Hadj-Chikh, L. Z., & Hazeltine, J. (1996). Caching and feeding decisions by *Sciurus carolinensis*: Responses to weevil-infested acorns. Journal of Mammalogy, 77, 305–314. doi:10.2307/1382802

45. Steele, M. A., Halkin, S. L., Smallwood, P. D., McKenna, T. J., Mitsopoulos, K., & Beam, M. (2008). Cache protection strategies of a scatter-hoarding rodent: do tree squirrels engage in behavioural deception? Animal Behaviour, 75, 705–714. doi:10.1016/j.anbehav.2007.07.026

46. Steele, M. A., & Koprowski, J. L. (2001). North American tree squirrels. Washington, DC: Smithsonian Institution Press.

47. Stevens, J. R., & Stephens, D. W. (2002). Food sharing: A model of manipulation by harassment. Behavioral Ecology, 13, 393–400. doi:10.1093/beheco/13.3.393

48. Stone, E. R., & Baker, M. C. (1989). The effects of conspecifics on food caching by black-capped chickadees. Condor, 91, 886–890.

49. Sun, S., & Zhang, H. (2013). Cache sites preferred by small rodents facilitate cache survival in a subtropical primary forest, central China. Wildlife Research, 40, 294–302.

50. Tamura, N., Hashimoto, Y., & Hayashi, F. (1999). Optimal distances for squirrels to transport and hoard walnuts. Animal Behaviour, 58, 635–642. doi:10.1006/anbe.1999.1163

51. Thompson, D. C., & Thompson, P. S. (1980). Food habits and caching behavior of urban grey squirrels. Canadian Journal of Zoology, 58, 701–710. doi:10.1139/z80-101

52. Vander Wall, S. B. (1990). Food Hoarding in Animals. Chicago: University of Chicago Press.

53. Vander Wall, S. B. (1995). The effects of seed value on the caching behavior of yellow pine chipmunks. Oikos, 74, 533–537.

54. Vander Wall, S. B. (2010). How plants manipulate the scatter-hoarding behaviour of seed-dispersing animals. Proceedings of the Royal Society B: Biological Sciences, 365, 989–997.

55. Vander Wall, S. B., & Jenkins, S. H. (2003). Reciprocal pilferage and the evolution of food-hoarding behavior. Behavioral Ecology, 14, 656–667. doi:10.1093/beheco/arg064

56. Waite, T. A., & Reeve, J. D. (1995). Source-use decisions by hoarding gray jays: Effects of local cache density and food value. Journal of Avian Biology, 26, 59–66.

